# Genomic Patterns of Parallel Divergence Across Demographically Heterogeneous Stickleback Populations in Eastern Canada

**DOI:** 10.1101/2025.04.18.649616

**Authors:** Alan García-Elfring, Antoine Paccard, Rowan D. H. Barrett

## Abstract

The threespine stickleback (*Gasterosteus aculeatus*) is a key model in evolutionary genetics, particularly for studies of parallel evolution, yet most genomic insights derive from populations on the west coast of North America and in Europe. Here, we use restriction site-associated DNA sequencing (RAD-seq) of pooled samples to examine genomic differentiation between marine and freshwater stickleback populations from Atlantic Canada. Our analyses reveal substantial heterogeneity in the extent and genomic distribution of marine–freshwater differentiation, with some freshwater populations showing strong divergence consistent with long-term isolation and drift, and others exhibiting patterns consistent with ongoing gene flow and admixture. Despite this demographic variation, we identify genomic regions that are repeatedly differentiated between marine and freshwater habitats, including loci near dopamine receptor genes (*Drd4a* and *Drd2l*). Gene ontology analyses of candidate regions show enrichment for functions related to nervous system development and dopamine receptor activity. Together, these results indicate that freshwater-associated genomic differentiation in Atlantic Canadian stickleback occurs across contrasting demographic contexts and suggest a potential role for neurological and behavioural pathways in adaptation to freshwater environments.

**Significance statement:** Threespine stickleback are a model system for studying parallel evolution, yet most genomic research has focused on Pacific and European populations. By examining previously understudied Atlantic Canadian populations, we identify genomic patterns consistent with parallel divergence across markedly heterogeneous demographic contexts, including populations shaped by strong drift as well as gene flow. We detect differentiation near dopamine receptor genes, pointing to a potential role for behavioural and hormonal pathways. Together, these results highlight the role of demographic context in shaping patterns of genomic differentiation associated with freshwater colonization.

## Introduction

The genetic basis of adaptation underpins the mechanisms of species diversification. A classic system for exploring these processes is the threespine stickleback (*Gasterosteus aculeatus*), native to the oceans of the Northern Hemisphere and an important species in ecological and evolutionary studies over the last century (Bertin 1925; Heuts 1947a, b; Moodie and Reimchen 1976; Nagel and Schluter 1998; Marques et al. 2018; Sanderson et al. 2026). Following the last glacial maximum (∼21 kya), retreating glaciers created new freshwater lakes and rivers into which marine stickleback populations dispersed and became isolated. Many of the nascent freshwater populations evolved similar low-plated morph phenotypes, particularly in western North America (Baumgartner and Bell 1984; Colosimo et al. 2005; Hohenlohe et al. 2010). Alleles that confer higher fitness in freshwater environments often persist at low frequencies in marine populations. Bidirectional gene flow is thought to maintain this standing genetic variation, facilitating the repeated colonization of freshwater habitats and the independent evolution of similar phenotypes (the “transporter hypothesis”; Schluter and Conte 2009). As a result, the threespine stickleback is widely regarded as a model system for studying parallel evolution.

However, even in the early stages of establishing stickleback as a model species, researchers noted that the striking parallel evolution observed among freshwater populations might be a geographically restricted phenomenon (Hagen and Moodie 1982), largely confined to western North America (eastern Pacific, e.g., Colosimo et al. 2005) and parts of Europe (e.g., Klepaker 1995). For instance, considering the armour phenotype, in several regions such as eastern North America, Northern Europe, the Baltic Sea, and eastern Asia, the completely plated morph—typical of marine environments—also predominates in local freshwater lakes (Münzing 1963; Penczak 1965; Hagen and Moodie 1982; Mäkinen et al. 2008; Raeymaekers et al. 2014; Ferchaud and Hansen 2016; Yamasaki et al. 2019). This limited phenotypic divergence between marine and freshwater populations has also been observed at the genetic level using allozymes (Rafiński et al. 1989). A large-scale genomic study by Fang et al. (2020a) confirmed that high levels of marine–freshwater (M-FW) genomic differentiation are primarily restricted to the eastern Pacific, whereas Atlantic populations show relatively low differentiation. In contrast to the strong divergence seen in British Columbia populations, many freshwater populations in Eastern Canada are dominated by the fully plated ecotype (Hagen and Moodie 1982; Haines 2023), although some exhibit distinct morphological variation (Scott et al. 2023). Despite these observations, genomic studies in this region remain limited (but see Fang et al. 2020a).

One potential explanation for the reduced extent of parallel evolution and M-FW genomic differentiation outside the eastern Pacific is the stochastic loss of freshwater-adapted alleles. Threespine stickleback originated in the Pacific Ocean, where they have persisted for approximately 26 million years (Matschiner et al. 2011; Betancur-R et al. 2015). It was only much more recently, during the late Pleistocene (36.9–346.5 thousand years ago), that they expanded their range into the western Pacific and the Atlantic basin (Orti et al. 1994; Fang et al. 2018; Fang et al. 2020b). As a result of this range expansion, Atlantic populations exhibit lower genetic diversity compared to their Pacific counterparts (Fang et al. 2020a). Notably, most Atlantic populations examined to date have been from Europe, leaving other regions, such as Atlantic Canada, relatively understudied.

Here, we investigate marine and freshwater threespine stickleback from eastern Canada, specifically from the provinces of Nova Scotia and Newfoundland, where fully plated phenotypes have been observed in freshwater habitats (Hagen and Moodie 1982; Haines 2023). Using genome-wide SNP data, we characterize M-FW genomic differentiation and test for signatures of parallel natural selection. Our goal is to gain insight into the evolutionary history and genetic basis of freshwater adaptation in stickleback populations from eastern North America.

## Results

### Genomic diversity, population structure, and differentiation among populations

We sampled 30 adult threespine stickleback from each of nine populations along the eastern Canadian Atlantic coast (N = 270 total), including four marine and five freshwater sites in Nova Scotia and Newfoundland (Figure 1; Table S1). Genome-wide estimates of nucleotide diversity (π), Watterson’s θ, and Tajima’s D varied among populations and habitats (Figure 2; Table S2). Marine populations like Antigonish Landing consistently showed high levels of genetic diversity (π ≈ 0.31–0.33) and positive Tajima’s D values, consistent with substantial standing variation. Freshwater populations exhibited greater heterogeneity. Blue Pond showed the lowest π and θ values and a near-zero to slightly negative Tajima’s D, indicating low genetic diversity consistent with strong drift/founder effects, followed by Lake Ainslie and Black River (Table S2). In contrast, Pomquet Lake and Pinchgut Lake showed among the highest π and Tajima’s D values observed, indicating an excess of variants at intermediate frequencies and potential gene flow.

**Figure 1.**
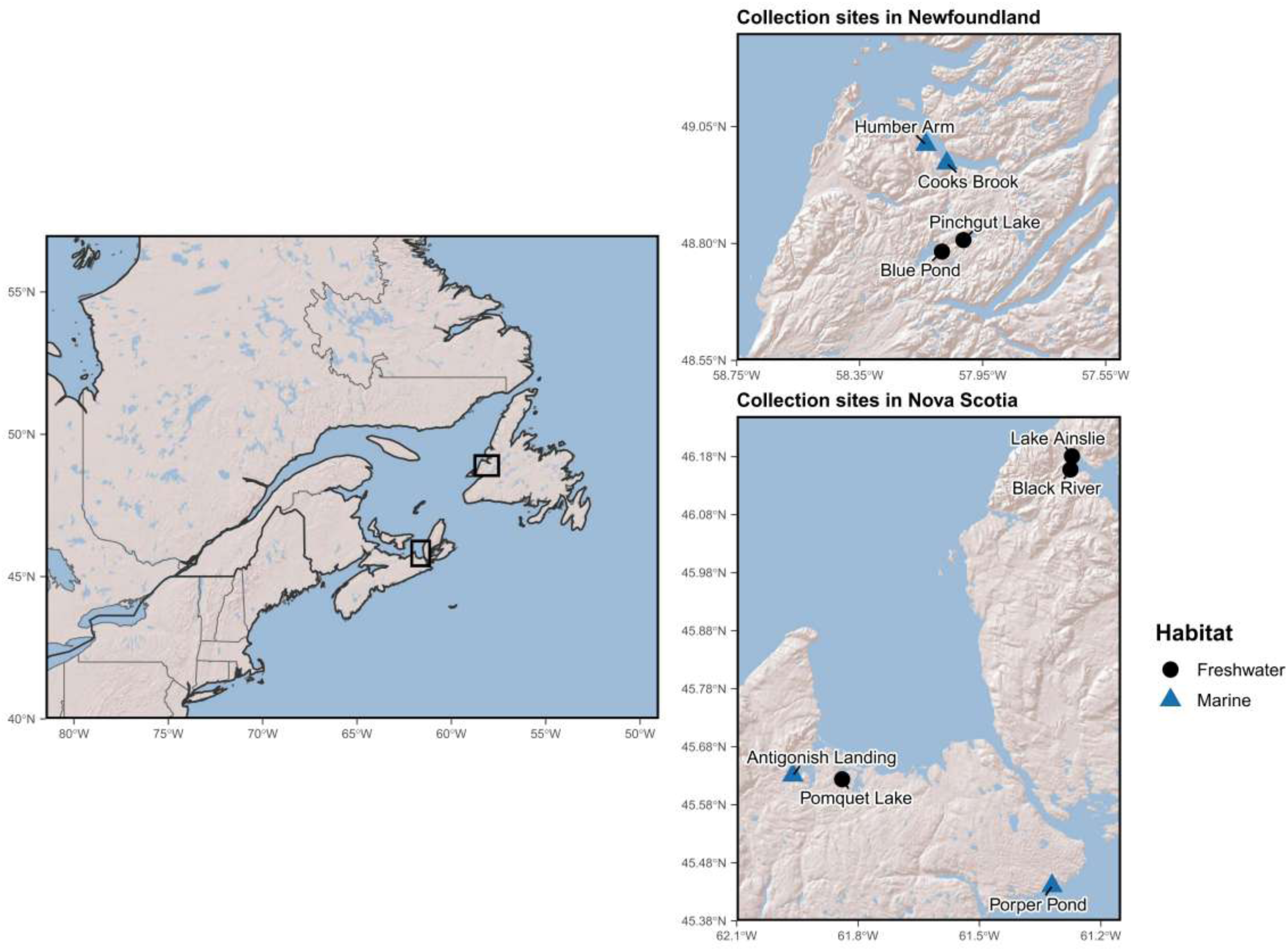
Sampling localities of threespine stickleback populations in eastern Canada. The left panel shows eastern North America, with sampled regions highlighted by black rectangles. Insets display collection sites in Newfoundland (upper right) and Nova Scotia (lower right). Freshwater populations are shown as black circles and marine populations as blue triangles.

**Figure 2.**
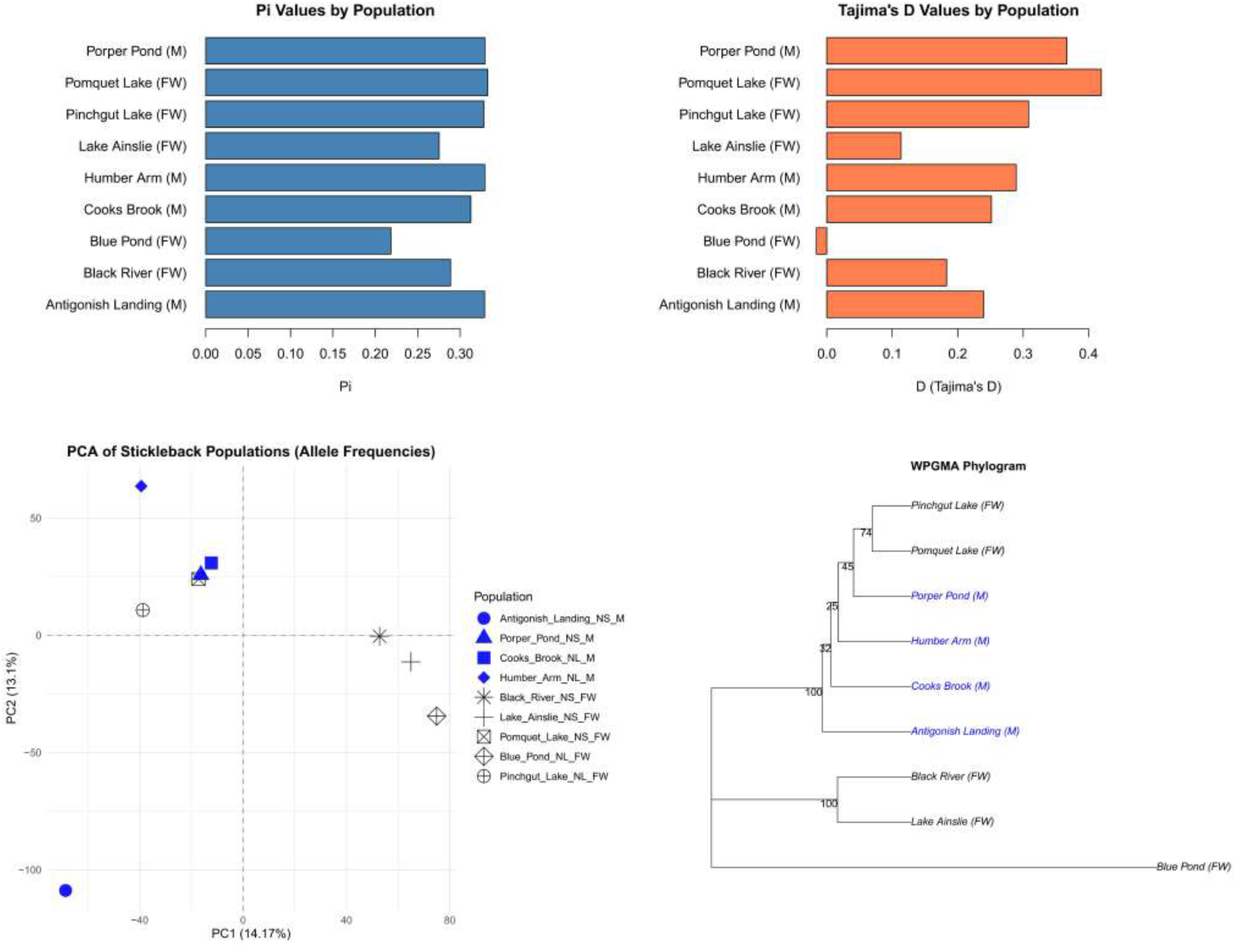
Population structure and summary population genetic statistics in Eastern Canadian threespine stickleback. (Top left) Principal component analysis (PCA) based on genome-wide SNP allele frequencies showing differentiation between marine (M) and freshwater (FW) populations; PC1 and PC2 explain 14.17% and 13.1% of the total variance, respectively. Points are coloured by habitat. (Top right) WPGMA phylogram constructed from genome-wide SNP allele-frequency distances illustrating relationships among populations; bootstrap support values are shown at nodes. (Bottom left) Nucleotide diversity (π) estimated for each population, summarizing within-population genetic variation. (Bottom right) Tajima’s D values by population, reflecting deviations in the allele frequency spectrum relative to neutral demographic equilibrium. Marine populations are indicated by (M) and freshwater populations by (FW).

To gain insight into population structure, we performed a principal component analysis (PCA) on allele frequencies (Figure 2, PCA). Marine and freshwater populations segregated into three main groups along PC1 (14.2% of variance explained) and PC2 (13.1%). Freshwater populations characterized by lower genetic diversity and lower Tajima’s D—Blue Pond, Lake Ainslie, and Black River—clustered together. In contrast, Pomquet Lake and Pinchgut Lake, freshwater populations with higher genetic diversity and Tajima’s D (putative gene flow from the ocean), clustered together with several marine populations (Porper Pond, Cook’s Brook, and Humber Arm), which extend further along PC1. The marine population Antigonish Landing formed a distinct cluster.

Hierarchical clustering using the WPGMA method grouped populations into clades that overall corroborate the PCA (Figure 2 phylogram): Lake Ainslie and Black River formed one sister group, with Blue Pond appearing as a more genetically divergent freshwater population. Pomquet Lake and Pinchgut Lake cluster together in a clade nested among the marine populations with which they overlap in the PCA, showing low bootstrap support at internal branches, suggesting weak genetic structure consistent with high connectivity and ongoing gene flow. To this group, Antigonish Landing forms a sister group with strong bootstrap support (100%), indicating genetic differentiation from the putatively admixed freshwater and marine clade.

We next examined population structure by visualizing median genome-wide F_ST_ across all 36 pairwise comparisons based on 18,582 SNPs as a heatmap (Figure 3; Table S3). Relative divergence varied widely, ranging from 0.0118 to 0.0824. Substantial genomic heterogeneity was observed among both marine–freshwater (M–FW) and freshwater–freshwater (FW–FW) comparisons (Figure 4; Figures S1, S2). Notably, both the lowest and highest levels of genomic differentiation occurred among freshwater populations: Pomquet Lake (Nova Scotia) and Pinchgut Lake (Newfoundland) were the least differentiated, whereas Lake Ainslie (Nova Scotia) and Blue Pond (Newfoundland) were the most differentiated.

**Figure .3.**
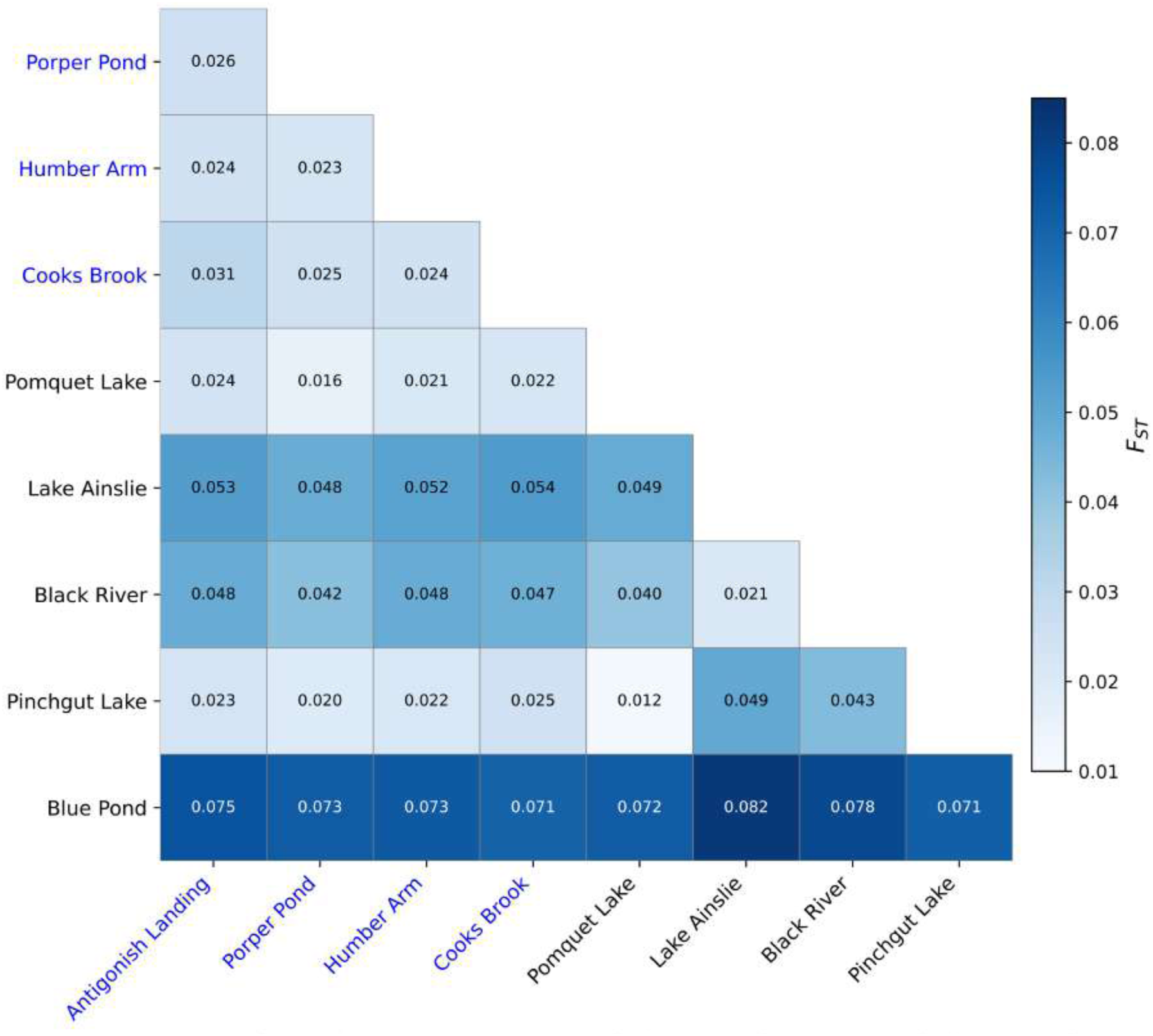
Heatmap of median pairwise F_ST_ among populations. Freshwater population are shown in black and marine populations in blue.

**Figure 4.**
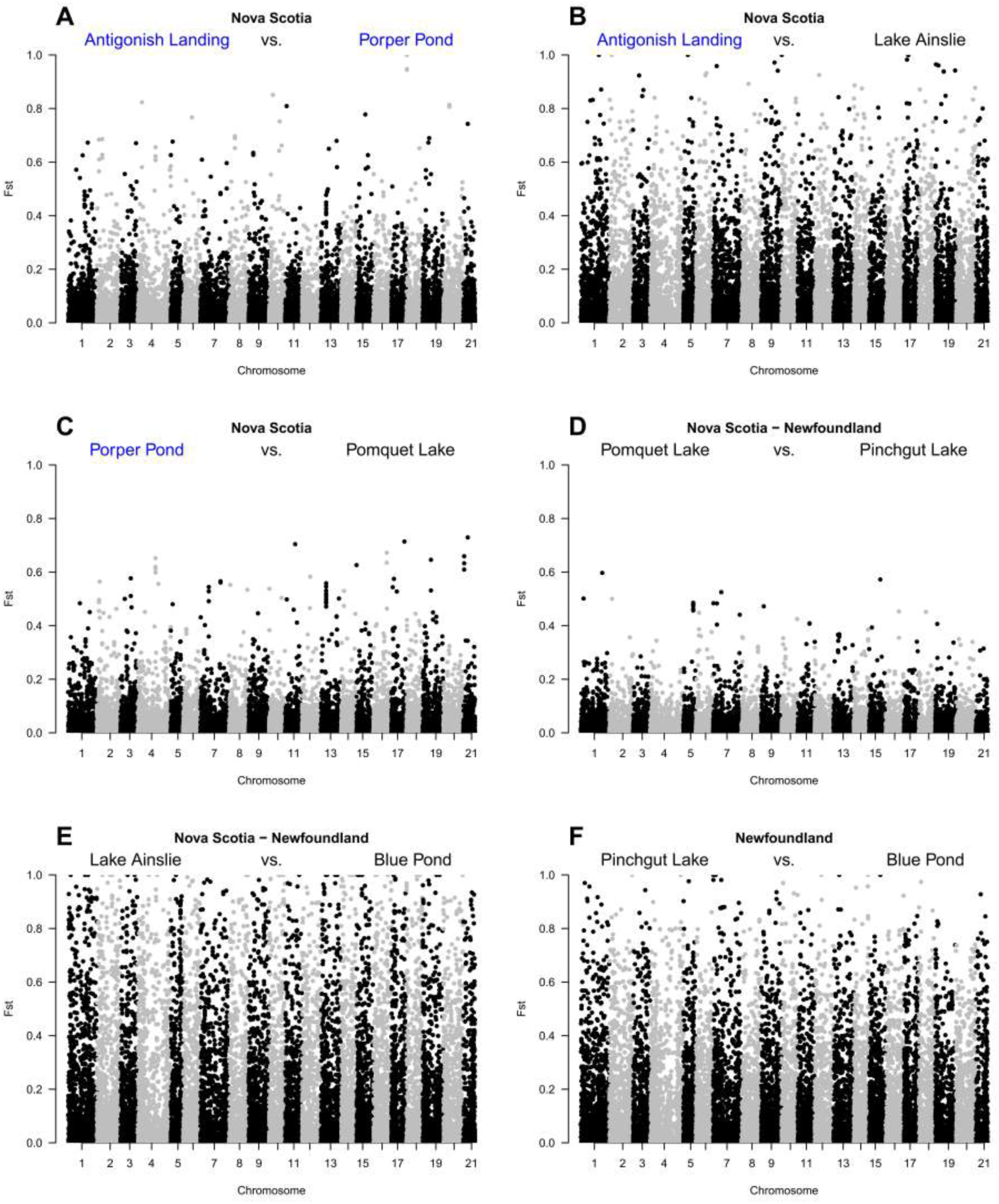
Manhattan plots of FST values highlighting heterogeneity among threespine stickleback populations from Eastern Canada. Freshwater populations are labelled in black and marine in blue.

Comparisons of relative (F_ST_) and absolute (d_XY_) divergence across 20 kb windows revealed a decoupling between these two measures (Figure 5). Across key population pairs (e.g., Figure 4), windows with high F_ST_ did not show correspondingly elevated d_XY_, indicating that extreme allele frequency divergence is not driven by increased sequence divergence between populations. Instead, high F_ST_ values occur in windows with low d_XY_, a pattern expected in populations that have experienced genetic drift and possess lower genetic diversity.

**Figure 5.**
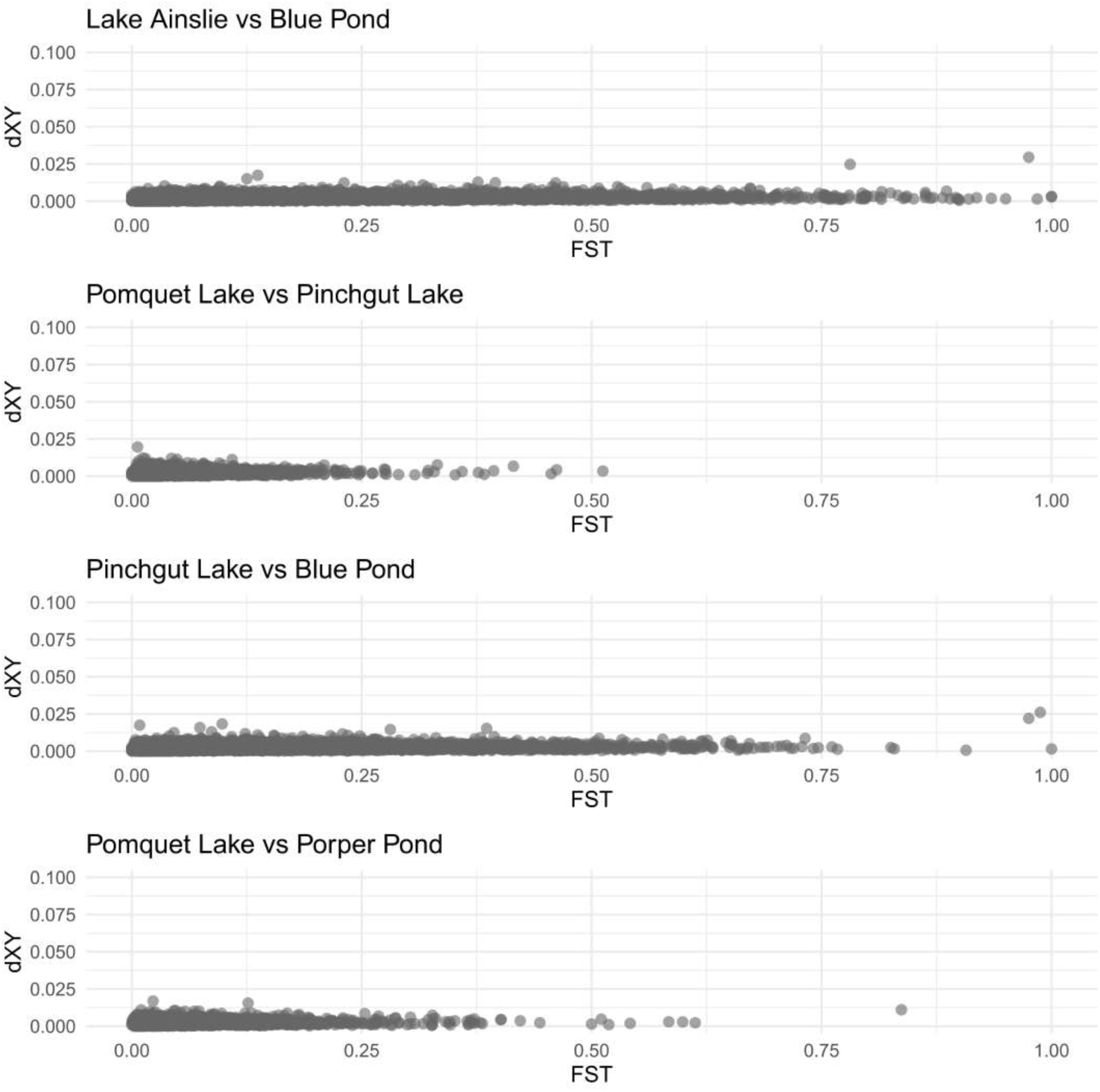
Comparison of absolute (dXY) and relative (FST) divergence among populations experiencing heterogeneous demographic histories (as inferred by estimates of Tajima’s D).

### SNPs showing repeated divergence between marine and freshwater populations

Excluding comparisons between populations from the same habitat type (M-M and FW-FW), we obtained 25,236 SNPs across the 20 possible M-FW comparisons within and across provinces. We used this SNP set to conduct an exploratory analysis to identify genomic regions showing repeated differentiation between marine and freshwater populations, with the aim of generating candidate loci potentially involved in parallel freshwater adaptation (Figure 6). We identified 2,100 SNPs as F_ST_ outliers in two or more M-FW comparisons within Nova Scotia (19 SNPs across all six comparisons). In Newfoundland, we identified 1,306 SNPs as F_ST_ outliers in two or more comparisons. Comparing interprovincial differentiation among 10 M-FW population comparisons revealed 3,422 SNPs as F_ST_ outliers in at least two comparisons, with three SNPs appearing in nine of the 10 comparisons (Figure 6). In the top 5% of the F_ST_ distribution, we identified 279 SNPs (referred to as ’F_ST_ outliers’), which were outliers in at least 6/20 M-FW comparisons, representing 1.1% of the SNP dataset.

**Figure 6.**
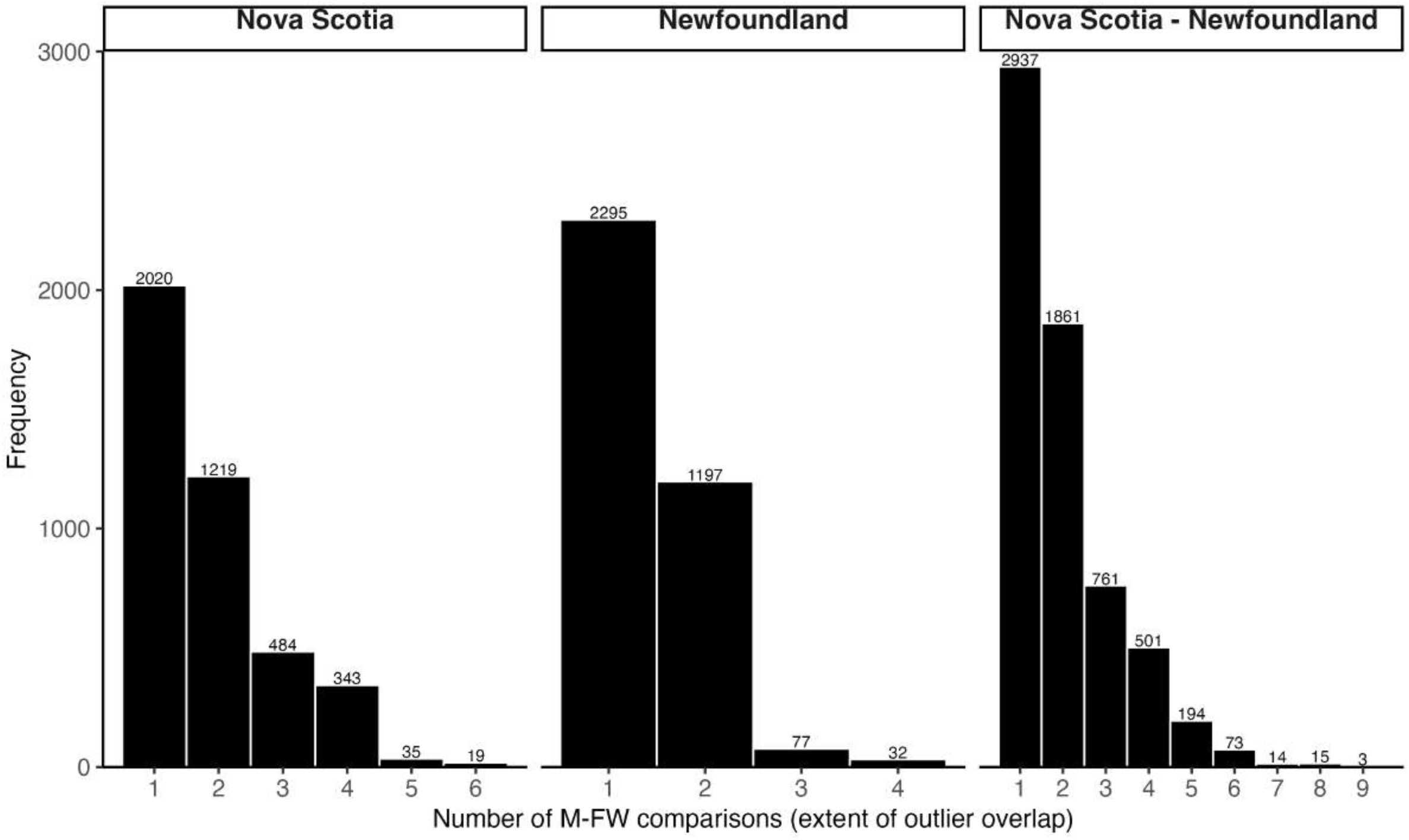
Overlap of FST outlier SNPs (top 5%) from marine–freshwater (M–FW) comparisons within and between regions. Bars show the number of SNPs shared among increasing numbers of comparisons, and numbers above each bar indicate the total SNPs detected in that overlap category.

We found that 180 of 279 F_ST_ outliers map within 5 kb of 133 genes (Table S4), with 68 genes containing at least one outlier (Table S5). Notably, two SNPs (T>C at VI: 16449327 and C>T at XX: 14009648) were in the top 5% of F_ST_ values in 19/20 M-FW comparisons (6/6 in Nova Scotia, 4/4 in Newfoundland, and 9/10 interprovincial comparisons; Figure 6). The latter SNP is located 920 bp downstream from a dopamine receptor gene, *Drd2l* (XX: 14002115-14008728). It is worth noting that these markers show lower coverage in some populations, which is known to bias levels of F_ST_. (VI: 16449327 coverage range: 5-21X; XX:14009648 coverage range: 5-35X). However, although both alleles (T and C) were present in marine populations at both loci (VI:16449327 T:C = 11:39; XX:14009648 T:C = 33:63), the C allele was almost entirely absent from freshwater populations. Across freshwater habitats, T:C coverage ratios were 38:2 at the chromosome VI locus and 55:1 at the chromosome XX locus. Notably, the only freshwater population in which the C (“marine”) allele was detected was Pomquet Lake.

To formally assess significance, we applied a Fisher’s Exact Test (FET), which identified 18 SNPs with significant allele frequency differences (FDR < 0.10) across the 10 marine–freshwater comparisons, each located within 5 kb of different genes (Table S6). Three of these genes also overlap with FST outlier genes (*dmap1*, *hnrnpm*, and *drd2l*). The armour-plating locus (IV:12811481) was not sampled in our RAD-seq dataset, with the closest SNP mapping 72.3 Kb downstream (IV:12883820).

#### Gene Ontology

Genes within 5 kb of FST outliers were enriched for biological processes related to adenylate cyclase-inhibiting dopamine receptor signalling and nervous system development (Table S7). Enriched molecular functions included neurotrophin binding and dopamine neurotransmitter receptor activity (Table S8).

## Discussion

In this study, we investigated marine and freshwater populations of threespine stickleback along Canada’s east coast to gain insights into their evolutionary history and the potential genetic basis of freshwater adaptation. Contrary to expectations of low genome-wide differentiation between marine and freshwater populations in the western Atlantic (Fang et al. 2020a), we found that differentiation among M-FW populations varied widely along the east coast of North America. This suggests a complex evolutionary history, likely involving multiple colonization events, distinct glacial refugia, or ongoing gene flow. Evidence of parallel evolution was observed as repeated allele frequency differences at SNP loci across several M-FW population comparisons. Notably, we identified a consistent signal of differentiation at one SNP near a dopamine receptor previously found to be differentially expressed between marine and freshwater populations (Di-Poi et al. 2016). Our results suggest that, even in the likely absence of key alleles for reduced armour in eastern Canadian populations, behavioural or endocrine adaptations may have facilitated freshwater persistence.

### Heterogeneous marine-freshwater differentiation in Eastern Canada

Current evidence suggests that threespine stickleback colonized the Atlantic during the late Pleistocene, as recently as ∼36.9 thousand years ago (Fang et al. 2018; Fang et al. 2020b). Repeated bottlenecks during this expansion may have led to the stochastic loss of some freshwater-adaptive alleles, such as the low-plated Eda allele (Fang et al. 2020a). Loci under selection in freshwater are expected to differentiate rapidly following colonization. Thus, the low M-FW differentiation observed in many Atlantic populations may reflect a lack of such adaptive alleles in their standing genetic variation, compared to Pacific populations.

Here, we show that while genome-wide M-FW differentiation is indeed low in some eastern North American comparisons, other comparisons show levels of differentiation comparable to those seen on the Pacific coast (Figure 4, Figure S2). Genome-wide π and θ were consistently higher in marine populations (π ≈ 0.31–0.33) than in freshwater populations, which exhibited greater heterogeneity. Freshwater populations with low π and θ, such as Blue Pond, Lake Ainslie, and Black River, showed near-zero or slightly negative Tajima’s D, consistent with reduced genetic diversity and neutral evolution, whereas Pomquet Lake and Pinchgut Lake showed higher π and positive Tajima’s D, suggesting an excess of intermediate-frequency variants possibly due to ongoing gene flow. This inference was supported by PCA, where Pomquet Lake and Pinchgut Lake clustered with marine populations rather than with other freshwater populations, like Lake Ainslie and Black River, indicating likely gene flow between these freshwater populations and nearby marine sources.

The highest levels of differentiation were observed between different freshwater populations. For example, two lakes in Newfoundland—Pinchgut Lake and Blue Pond—are only 5 km apart but differ substantially in their genetic divergence from marine populations (Figure 4). Blue Pond is a small, rain-fed lake without input or output streams, making gene flow from neighbouring populations unlikely. Such barriers to gene flow likely promoted divergence through genetic drift. Comparisons of F_ST_ and absolute divergence (d_XY_) revealed a decoupling of relative and absolute divergence (Figure 5), with high F_ST_ windows showing low d_XY_, consistent with drift in low-diversity freshwater populations.

Two recent preprints provide important context for interpreting the structure of the marine populations included in our study. Samuk et al. (2025) and Sumarli et al. (2025) analyzed RADseq and whole-genome data, respectively, from several of the same Nova Scotia marine sites sampled here. They found two genetically distinct marine ecotypes — “white” and “common” sticklebacks — co-occur in sympatry across the region, with ongoing gene flow and very low but detectable genomic divergence (FST ≈ 0.01). The common marine stickleback in Nova Scotia could be separated into two genetic clusters, “mainland” and “Bras d’Or”. Despite this shallow divergence, the white ecotype exhibits consistent differences in morphology, reproductive traits, and behaviour, including smaller body size, distinctive coloration, spine length, altered reproductive investment, divergent nesting substrate and location, modified parental care strategies, and distinct courtship displays (Samuk et al. 2025). In contrast, no trophic or ecological differentiation between ecotypes was observed.

Two of our marine sampling sites were also studied by Samuk et al. (2025). In their investigation, Antigonish Landing (western Nova Scotia) consisted almost entirely of individuals from the “common mainland” genetic cluster, whereas Porper Pond (eastern Nova Scotia) was composed almost entirely of individuals belonging to the white ecotype genetic cluster. Fish with ancestry belonging to the common mainland, common Bras d’Or, and white ecotype predominate in western, northern, and eastern Nova Scotia, respectively. We did not observe any obvious phenotypic differentiation among our marine populations sampled in Nova Scotia and Newfoundland, most notably, none exhibited the distinct colouration of the white stickleback ecotype. Given the low divergence and recent estimated split time (∼1,000 years), white and common sticklebacks share the vast majority of their genetic variation. Thus, even if a few undetected white sticklebacks individuals were included in our sample, their allele-frequency contribution would be unlikely to account for the localized, parallel allele-frequency shifts observed near dopamine receptor genes.

Our population structure analyses are concordant with the structure reported by Samuk et al. (2025). Principal component analysis (PCA) clearly shows Antigonish Landing and Porper Pond as distinct genetic clusters. Notably, Porper Pond clusters closely with freshwater populations Pomquet Lake and Pinchgut Lake, consistent with ongoing gene flow or admixture. One possible interpretation is that white sticklebacks represent a hybrid population (Marques et al. 2019). It is known that freshwater stickleback are smaller than marine (Haglund et al. 1990; Aguirre et al. 2022), can display lighter colouration (Miller et al. 2007;), and show an array of different behaviours (McPhail and Hay 1983; Di-Poi et al. 2014). However, neither Samuk et al. (2025) nor Sumarli et al. (2025) included freshwater populations in their analyses, limiting direct inference about the contribution of freshwater–marine connectivity to this structure.

Multiple colonization events in eastern Canada, similar to the patterns observed in the Japanese Archipelago (Kakioka et al. 2020) and Europe (Mäkinen et al. 2008) could have led to heterogeneity in genetic distances among freshwater populations. However, if freshwater populations had originated from separate sources or refugia (e.g., Mäkinen and Merilä 2008), they would have accumulated sequence divergence over time, resulting in high d_XY_. Instead, the low d_XY_ coupled with high F_ST_ indicates recent population divergence with minimal absolute sequence differentiation, consistent with colonization from a single or shared marine source followed by rapid drift-driven divergence in isolated freshwater lakes. Pinchgut Lake is connected to other lakes through Georges Lake, with a stream that extends to the ocean, while Pomquet Lake is a few kilometres (<4 km) from the ocean and likely experiences ongoing gene flow via Taylor Creek. In contrast, Blue Pond is a small, rain-fed lake without streams. This geographic variation in connectivity explains the heterogeneous population structure better than multiple independent colonization events from divergent sources.

### Repeated differentiation near dopamine receptor genes

Adaptation to similar environments via re-use of standing genetic variation is considered a primary mechanism underlying parallel evolution (Schluter and Conte 2009; Schlötterer 2023). The low-plated Eda allele, though present at low frequencies in marine populations, confers a fitness advantage in freshwater (Marchinko and Schluter 2007; Barrett et al. 2008; Le Rouzic et al. 2011). Its widespread loss in the Atlantic gene pool may explain the absence of parallel evolution for the low-plated phenotype in eastern Canadian lakes. However, *Eda* is not the only gene involved in freshwater adaptation; genes involved in osmoregulation, such as Na⁺/K⁺-ATPase and Na⁺/K⁺/2Cl⁻ cotransporters, have been repeatedly implicated across global stickleback populations (Deagle et al. 2013; Divino et al. 2016; McCairns and Bernatchez 2010; Garcia-Elfring et al. 2021).

Consistent with previous observations and the predicted loss of the freshwater Eda allele, we did not find signs of molecular evolution near the Eda locus in our populations (Hagen and Moodie 1982; Fang et al. 2020a), although our nearest marker was more than 70 Kb away. Despite overall low differentiation, we identified signals of parallel evolution at 1.1% of SNPs, which showed repeated marine–freshwater divergence both within and across Nova Scotia and Newfoundland. Notably, this included SNPs located near dopamine receptor genes *Drd4a* and *Drd2l*, and candidate genes were enriched for functions related to dopamine receptor activity and nervous system development. Strikingly, the ’C’ alleles of two parallel outlier SNPs were almost entirely missing in freshwater populations but remained polymorphic in marine populations, consistent with selection favouring one allele, acting against the other, or both. We provide primers for each locus that were designed *in silico* (Untergasser et al. 2012); these primers have not yet been experimentally tested (Table S9).

While individual variation in endocrine function remains less studied than other physiological traits in evolutionary biology, it holds strong potential for linking genetics, physiology, and ecology (Williams 2008). In zebrafish (*Danio rerio*), expression levels of dopamine receptors *Drd2c* and *Drd3* have been associated with individual aggression, particularly in dominant versus subordinate males (Filby et al. 2010). In stickleback, exposure to predators has been shown to reduce *Drd4a* expression (Abbey-Lee et al. 2018), and marine and freshwater populations exhibit marked differences in behaviours such as sociability, aggression, and activity (Di-Poi et al. 2014). At the population level, Di-Poi et al (2016) showed evolutionary changes in physiological regulatory networks often target receptor genes—such as *Drd2*—rather than their ligands or neurotransmitters. In a separate study using stickleback from a Nova Scotia lineage (Blouw and Hagen 1990), *Drd2* expression was associated with male behavioural trade-offs between courtship and territorial defence (Barbasch et al. 2023).

The dopaminergic system also plays a key role in osmoregulation through inhibition of prolactin, a hormone critical for ion retention and water balance in freshwater environments (Liu et al. 2006; Yamamoto and Vernier 2011; Mancera and McCormick 2019). Prolactin mRNA levels correlate with local ion concentrations in natural populations of black-chinned tilapia (*Sarotherodon melanotheron*) adapted to different salinities (Tine et al. 2007). In the euryhaline fish *Scatophagus argus*, renal dopaminergic signalling via *Drd1* inhibits Na⁺/K⁺-ATPase under hypo-osmotic stress (Su et al. 2016). In stickleback, prolactin is associated with freshwater migration and salinity tolerance (Lam and Hoar 1967; Ishikawa et al. 2016; Pavlova et al. 2020; <u>Taug</u>bøl et al. 2022). Taken together with the population genetic patterns described above, these results highlight dopaminergic signalling as a plausible axis of freshwater adaptation in eastern Canadian stickleback. However, because our dataset is pooled RAD-seq and we cannot estimate local LD decay, SNP-to-gene proximity should be treated as suggestive, and these dopamine receptor loci are best viewed as candidates motivating follow-up work. Future studies combining whole-genome data with functional assays and expression profiling will be required to test whether variation at specific loci contributes to adaptation in these systems.

## Conclusion

Our study revealed heterogeneous genomic differentiation between marine and freshwater threespine stickleback populations in eastern Canada. Some lakes, in particular Blue Pond, appear to represent drifted “island” populations with limited gene flow, whereas others, like Pomquet Lake, maintain higher diversity through ongoing connectivity with marine populations. Despite this difference in demographic history, we detected evidence of parallel evolution near dopamine receptor genes, where specific alleles are largely absent in freshwater but polymorphic in marine populations. Our results suggest that parallel evolution in eastern Canada may involve selection on behavioural (or osmoregulatory) phenotypes, shaped by both drift in isolated populations and on-going selection in connected populations, though further work is needed to test this hypothesis.

## Methods

### Field sampling and DNA extraction

We collected 30 adult threespine stickleback (> 30 mm standard length) from nine populations in eastern (Atlantic) Canada, comprising four marine and five freshwater populations. Fish were sampled using minnow traps and beach seines during June and July 2014 in Nova Scotia and Newfoundland, Canada (Figure 1; Table S1). These samples form the basis of the RAD-seq dataset analyzed in this study.

In Nova Scotia, freshwater populations were sampled from two lakes (Pomquet Lake and Lake Ainslie) and one stream (Black River), while marine samples were obtained from Antigonish Landing and Porper Pond. In Newfoundland, freshwater populations were sampled from Pinchgut Lake and Blue Pond, and marine populations from Cooks Brook and Humber Arm (Figure 1; Table S1). Genomic DNA was extracted from all individuals using a standard phenol–chloroform protocol.

### DNA sequencing of pooled samples

DNA concentration was quantified using a Picogreen® ds DNA assay (Thermo Fisher Scientific, Waltham, USA) and an Infinite® 200 Nanoquant (Tecan Group Ltd. Männedorf, Switzerland). DNA concentrations were normalized across individuals and pooled by sampling location, resulting in nine pooled samples, each consisting of 30 individuals.

RAD-seq libraries were prepared following the protocols of Paccard et al. (2020). Briefly, pooled samples were digested with NlaIII and MluCI, ligated to unique barcoded adapters, size-selected for ∼400 bp fragments, enriched for biotin-tagged fragments using streptavidin beads, and PCR-amplified. Cleanup steps were performed using AMPure XP beads. Libraries were sent for sequencing at the McGill University and Genome Quebec Innovation Center (Montreal, Canada) on one lane of Illumina HiSeq2500 platform, generating 125bp paired-end reads.

### Read processing and variant mapping

RAD-seq data was demultiplexed into pool-specific read sets using the *process_radtags* module of Stacks (Catchen et al. 2013), based on the barcode sequences listed in Table S1. We used the *trim-fastq*.*pl* script of *Popoolation* (Kofler et al. 2011a) to process raw reads, filtering based on read quality (--quality-threshold 20) and length (--min-length 50). We then mapped the processed reads to the stickleback reference genome (BROAD S1) with the program *Bowtie2* (Langmead and Salzberg 2012), using the --end-to-end mapping option. We used *SAMtools* (Li et al. 2009) to convert the output SAM files to BAM format and subsequently removed reads with mapping quality below 20 (samtools view -q 20). We generated an mpileup file (samtools mpileup -B) and converted the mpileup file to the synchronized (sync) format using *Popoolation2* (Kofler et al. 2011b), which was used for downstream analyses.

### Population genetic parameters, population structure, and allele frequency differentiation

#### Population genetic summary statistics

We estimated population genetic parameters, including nucleotide diversity (π), Watterson’s θ, and Tajima’s D, using the *variance-sliding.pl* script from PoPoolation2 (Kofler et al. 2011b).

#### Allele frequency estimation and PCA

To visualize population structure, we performed a principal component analysis (PCA) based on SNP-specific allele frequencies at loci shared across populations, which primarily reflect shared (ancestral) standing variation rather than population-specific rare variants. For each population, allele frequencies were obtained from cleaned and sorted BAM files which were processed using bcftools to generate downstream VCFs. Genome-wide variant calling was performed with bcftools mpileup and bcftools call. From these VCFs, sites were subset to a predefined panel of 18,582 SNP positions shared across populations (datasets defined below).

Allele depths (AD) for reference and alternate alleles were obtained at each SNP using bcftools query, and allele frequencies were computed as the fraction of reads supporting the alternate allele over the total read depth. Population-specific allele frequency files were then combined into a single matrix with SNPs as rows and populations as columns. Before performing principal component analysis (PCA), missing allele frequencies—caused when reads were present but alternate alleles were not confidently called by bcftools in the VCF AD field for certain SNP–population combinations—were imputed using the mean allele frequency for that SNP across populations. Unlike bcftools, Popoolation2 does not require confident genotype calls, only read counts from the sync file. SNPs with zero variance across populations were excluded. PCA was performed in R using *prcomp* on the centered and scaled allele frequency matrix, treating populations as observations. Population clustering was visualized using the first two principal components, with populations labeled and coloured according to habitat type (marine vs. freshwater).

We estimated relative genetic differentiation among populations using F_ST_, as defined by Hartl and Clark (1997), with the *fst-sliding.pl* script from PoPoolation2 (Kofler et al. 2011b). We visualized genetic relationships among marine and freshwater populations by constructing a WPGMA phylogram based on median of these F_ST_ values. A distance matrix was generated from genome-wide SNP data, and clustering was performed in R using the hclust(method = “mcquitty”) function. The resulting dendrogram was visualized as an unrooted phylogram using phangorn::plotBS(), with branch lengths scaled to genetic distance. Bootstrap support values were calculated from 10,000 replicates, and only values ≥10% are shown. The tree was explicitly unrooted to avoid assumptions about ancestral directionality.

Low sequencing coverage can bias F_ST_ estimates through sampling variance in allele frequencies. We therefore additionally assessed allele frequency differences using Fisher’s Exact Test (FET) implemented in *fisher-test.pl of* Popoolation2. Analyses were performed using the following parameters: --min-count 2, --min-coverage 5, --max-coverage 500, --min-covered-fraction 0, --window-size 1, --step-size 1, --pool-size 30:30:30:30:30:30:30:30:30, and --suppress-noninformative. Only genomic regions assembled at the chromosome level were analyzed; unplaced scaffolds were excluded. Also, because isolated populations experiencing strong genetic drift can exhibit elevated F_ST_ values as a consequence of reduced within-population diversity, we calculated absolute sequence divergence (d_XY_). Estimates of d_XY_, which is equivalent to between-population nucleotide diversity, were obtained using grenedalf (Czech et al. 2024), a framework designed, like Popoolation2, for Pool-seq data.

We next sought to identify variants putatively associated with parallel selection. For each marine–freshwater (M–FW) population comparison, we identified SNPs in the upper 5% of the F_ST_ distribution (e.g. Yan et al. 2025). This threshold was applied heuristically to retain a broad range of allele frequency differences across population pairs with heterogeneous demographic histories.To identify loci with consistent differentiation (i.e., putative parallelism), we defined three sets of SNPs in the top 5% across at least two M-FW comparisons: within Nova Scotia, within Newfoundland, and across both provinces. SNPs shared among all three sets were designated as “F_ST_ outliers.”

To complement the F_ST_ approach, we performed a genome-wide FET scan across 10 intraprovincial M-FW comparisons to identify SNPs showing consistent allele frequency shifts. To control for multiple testing, raw p-values were converted to q-values using the qvalue package (Storey et al. 2003), and SNPs with a false discovery rate (FDR) below 10% were retained. This FDR threshold reflects the expected proportion of false positives among significant results.

### Identification of candidate genes and analysis of molecular function

Outlier SNPs are expected to be located within genes potentially under parallel selection or, more commonly, be located near them, given that we used a reduced representation approach. To identify such genes, we applied a custom bash script to map outlier loci to or near protein-coding genes in the reference genome. Our search was limited to 14,252 protein-coding gene annotations, filtered by the attributes “ID=gene” and “biotype=protein_coding.” Given that our data represent a reduced portion of the genome, the causal mutation is unlikely to be directly sampled and is more likely in linkage disequilibrium (LD) with a nearby SNP. Because our dataset is pooled RAD-seq, we did not estimate local LD decay. To account for this, we examined genes either associated with outlier loci or located within a heuristically chosen window of 5 kb around them (e.g., Nichols et al. 2016; Chen et al. 2018). To investigate the traits putatively under selection, we analyzed the candidate genes for enrichment of molecular functions using ShinyGO 0.77 (Ge et al. 2020), which links gene lists to functional categories, such as gene ontology (GO), based on annotation databases from Ensembl and STRING-db. We assessed enrichment for GO terms in the categories of biological process and molecular function, using a false discovery rate (FDR) threshold of 10% and reporting the 20 most significant results. We compared our list of candidate genes to a background set of 14,252 genes from which outliers were sampled, using the ’best matching species’ option for further analysis.

### SNP and windowed datasets

Our analyses used two SNP datasets and one dataset based on 20 kb windows. The first SNP dataset included 18,582 variants (average minimum coverage across populations: 56.4X) shared across all nine populations and present in all 36 pairwise population comparisons, including within- and between-habitat types (M-M, FW-FW, and M-FW). This dataset was used to estimate population genetic parameters (π, θ, D), calculate F_ST_ across all 36 comparisons (e.g., Manhattan and heat plots), perform PCA, and conduct FET scans. For analyses of marine–freshwater parallelism, which excluded M-M and FW-FW comparisons, we used a second dataset of 25,236 SNPs (average minimum coverage among populations: 54.5X) informative in the 20 relevant pairwise comparisons.

When examining absolute divergence, we employed a 20 kb windowed approach, calculating F_ST_ with Popoolation2 and d_XY_ with grenedalf. We used Popoolation2 to set the parameters. Because reduced-representation data results in many 20 Kb windows with zero coverage, we retained informative windows by setting the --min-covered-fraction parameter (the minimum fraction of a window with coverage between 5-200X across all populations) to 0.0001, based on the median and mean distances between variants in the 18,582-SNP dataset (Mean distance: 21,417.65 bp; Median distance: 3,977.00 bp; Std deviation: 40,189.06 bp; Max distance: 1,385,936 bp). Only windows containing at least three SNPs were included in the analysis. The 0.0001 threshold ensured that all 20 Kb windows have at least some SNPs retained, while the ≥3-SNP filter adds statistical robustness by averaging each window’s estimate over multiple data points. This filtering resulted in a total of 7,160 F_ST_ windows from Popoolation2 (average minimum coverage at each SNP in the window: 24.5X). We subsetted the same 7,160 windows from grenedalf to directly compare F_ST_ to d_XY_ estimates.

## Acknowledgments

This work was supported by a Discovery Grant from the Natural Sciences and Engineering Research Council of Canada (NSERC) awarded to R.B., as well as funding from the Canada Research Chairs (CRC) program. We thank Mathilde Salamon for her assistance and valuable contributions throughout the project as well as Dieta Hanson and the Mullaney family for their assistance with field sampling.

## Data availability

All raw sequencing data generated in this study have been deposited in the NCBI Sequence Read Archive (SRA) under a BioProject accession PRJNA1406969.

---

**Figure S1.**
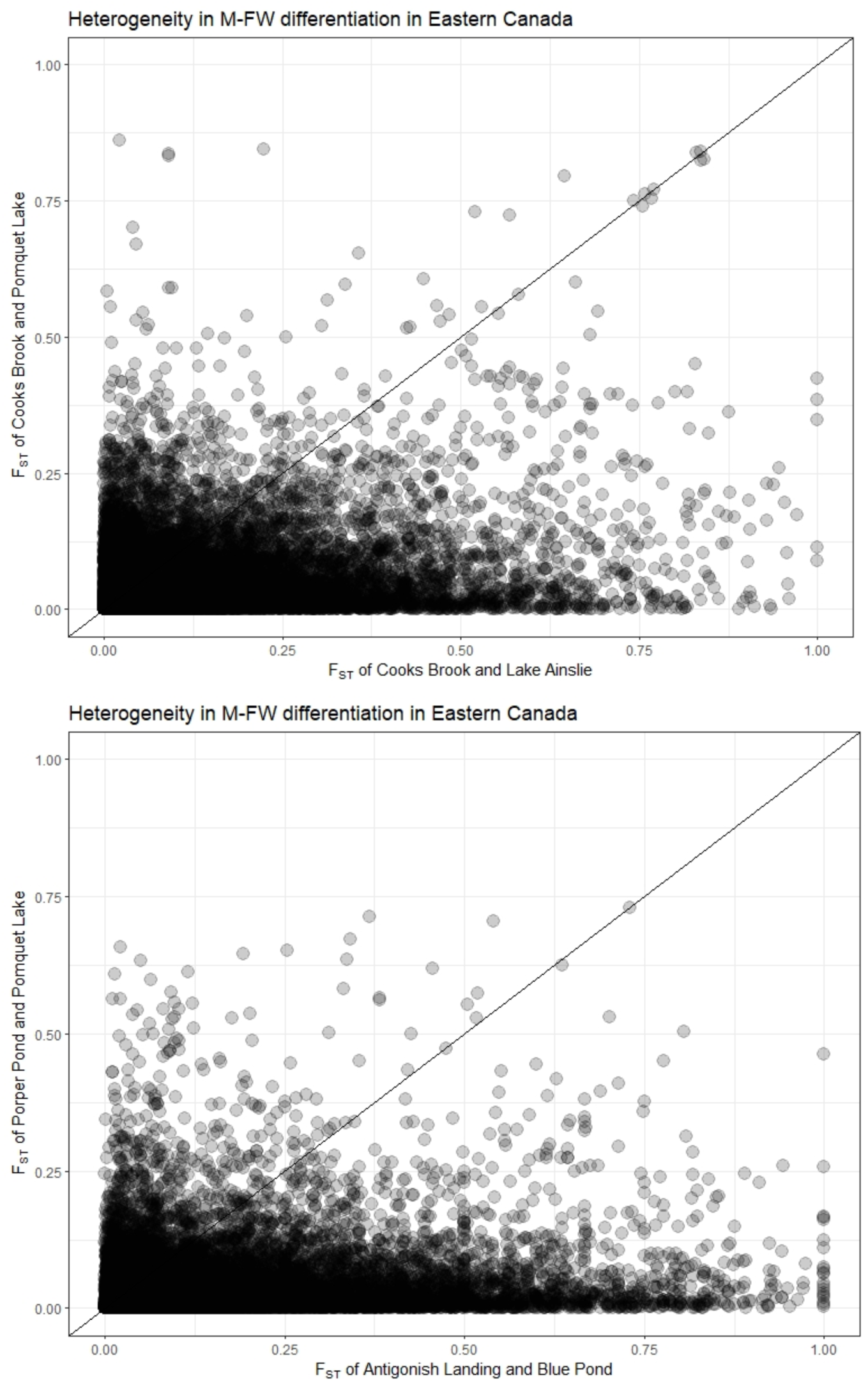
Heterogeneity of M-FW differentiation in Eastern Canada (Newfoundland and Nova Scotia).

**Figure S2.**
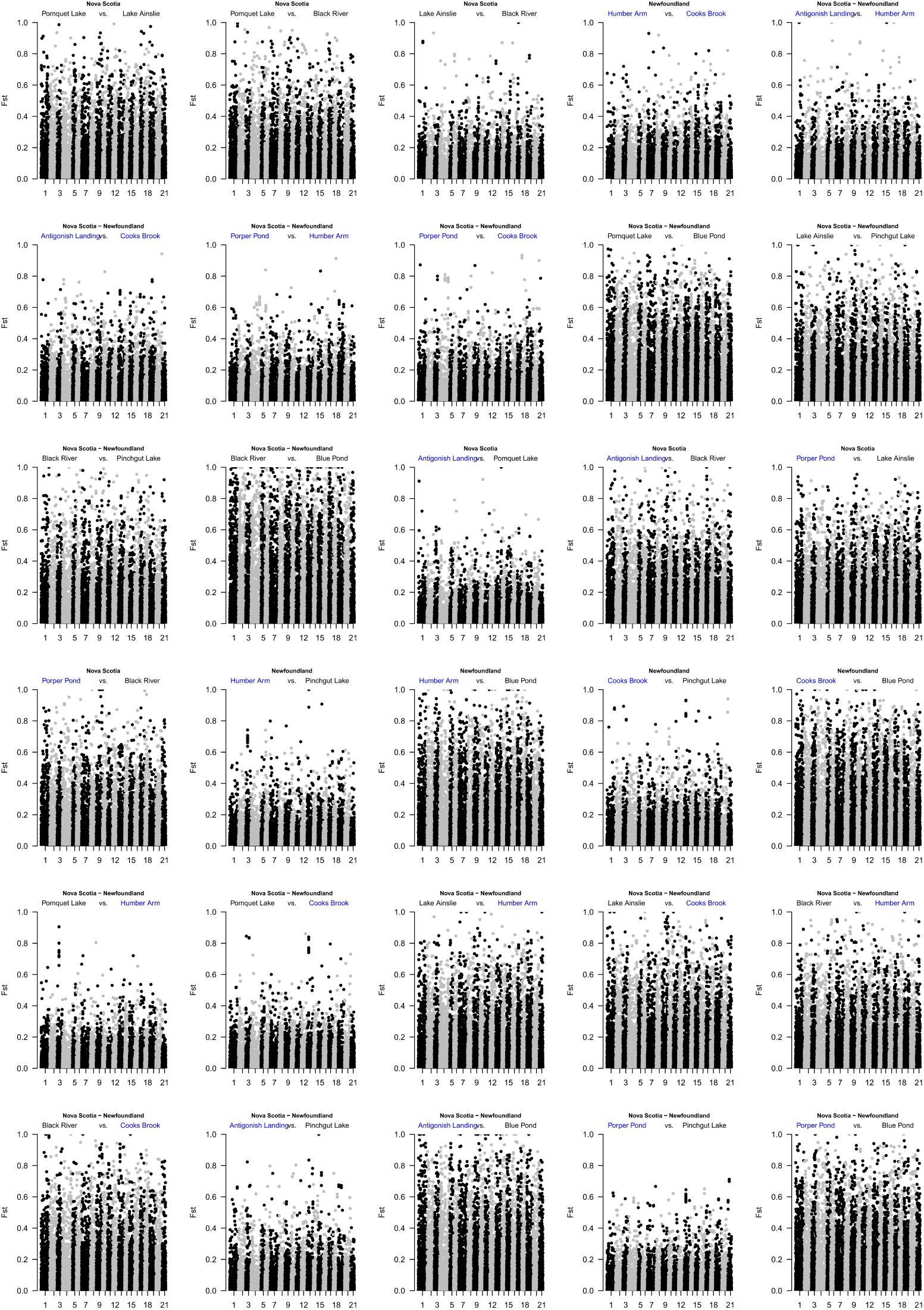
Manhattan plots of genome-wide differentiation across marine-marine, marine-freshwater, and freshwater-freshwater population comparisons within, and across, Nova Scotia and Newfoundland.

